# In exogenous attention, time is the clue: brain and heart interactions to survive threatening stimuli

**DOI:** 10.1101/2020.11.17.386334

**Authors:** Elisabeth Ruiz-Padial, Francisco Mercado

## Abstract

The capture of exogenous attention by negative stimuli has been interpreted as adaptive for survival in a diverse and changing environment. In the present paper, we investigate the neural responses towards two discrete negative emotions with different biological meanings, disgust and fear, and its potential relationships with heart rate variability (HRV) as an index of emotional regulation. With that aim, 30 participants performed a digit categorization task while fear, disgust and neutral distractor pictures were presented. Resting HRV at baseline, behavioral responses, and event-related potentials were recorded. Whereas P1 amplitudes were highest to fear distractors, the disgust stimulation led to augmented P2 amplitudes compared to the rest of distractors. Interestingly, increased N2 amplitudes were also found to disgust distractors, but only in high HRV participants. Neural source estimation data point to the involvement of the insula in this exogenous attentional response to disgust. Additionally, disgust distractors provoked longer reaction times than fear and neutral distractors in the high HRV group. Present findings are interpreted in evolutionary terms suggesting that exogenous attention is captured by negative stimuli following a different time course for fear and disgust. Possible HRV influences on neural mechanisms underlying exogenous attention are discussed considering the potential important role of this variable in emotional regulation processes.

## Introduction

To process the great amount of sensorial stimulation that continuously surrounds the individual a fast and precise brain selection is required in order to identify relevant signals from the environment such as those are important for survival to face them with an appropriate action (e.g.,[1]). Growing experimental evidence coming from both electrophysiological [2-7] and behavioral studies [8-11] has consistently shown that threatening information is capable to efficiently attract attentional resources in a rapid and automatic way (i.e., exogenous attention) when these stimuli appeared as distractors in a given visual task (for a review see [12]).

A new line of evidence has focused its efforts on exploring exogenous attentional responses to each type of negative-arousing events (i.e., fearful and disgusting) considering them as separate categories due to the subjective feelings related to each of them, their biological meaning, and even their neural substrate and effect on the peripheral nervous system are segregated [13-15]. Indeed, exogenous attention to affective events triggers fast neural responses reflected by enhanced amplitudes to threatening distractors in event-related potential (ERP components within the first 200 ms (P1, P2 and N2) from the stimulus onset [9]. In this sense, previous ERP studies have found neural indices (P2 wave) describing that disgust distractors are more efficient at attracting automatic attention than fearful distractors [14]. This segregated pattern between disgust and fear was also found when neural activation within visual regions (i.e., cuneus) was measured. More recently, Zhang and colleagues [16] using a dot-probe paradigm showed a clear attentional modulation of the N1 component in response to disgust stimuli being interpreted as a rapid suppression of attention to diminish risk of contamination.

With respect to peripheral physiological responses, research has consistently found that disgust produces greater heart rate (HR) deceleration compared to fear and neutral stimuli [17-20]. In fact, it has been proposed that this greater HR deceleration to disgusting stimuli would be specifically associated with the involvement of parasympathetic activation [17]. We have previously shown that startle and HR responses during fear and disgust pictures are modulated by vagally mediated heart rate variability (HRV). Participants with high HRV showed the expected startle magnitude increase to unpleasant foreground while the group with low HRV did not [21-22], and deeper cardiac deceleration to disgusting compared to fear-evoking or neutral distractors [23].

The importance of the bidirectional interactions between the heart and brain has been known for over 100 years [24]. HRV has been proposed as an index of the dynamics of brain–heart connections [25]. Specifically, vagal regulation of the heart indicates highly functional prefrontal inhibition of subcortical structures since the neural circuits bidirectionally connecting the prefrontal cortex with subcortical structures are linked to the heart via the vagus nerve [26-28]. High HRV has been associated with better affective and attentional regulation as indicated by, for example, larger orienting responses but faster habituation to nonthreatening stimuli [29] and adaptive emotional startle modulation [30]. However, low HRV has been related to hyperactive subcortical activity, which results in poor self-regulatory functions, such as the failure to recognize safety cues or to habituate to novel, neutral stimuli [26, 29, 31-32].

Closely connected to the present study, HRV has been recently related to the capture of automatic attention by emotional stimuli. Park et al [33] asked their participants to detect a target letter among a string of letters superimposed on either fearful or neutral distractor faces. Their results showed that neutral and fearful faces interfered with the performance of the primary task in the low HRV group whereas in the high HRV group the interference was observed only to the fearful faces. Park et al [34] showed faster attentional engagement to fearful faces in low HRV than in high HRV participants what was considered dysfunctional. These findings together suggest that only high HRV participants showed an adaptive response.

Despite ERPs are considered as a suitable tool to explore rapid neural responses occurring at early attentional processing stages, such as exogenous attention, its study to characterize segregated neural substrates related to fear and disgust as a function of individual differences in resting HRV has not been explored. Furthermore, the use of mixed experimental paradigms in studies on exogenous attention to disgust and fear stimuli has led to obtain inconclusive outcomes that hinder the possibility to shed light on the underlying mechanisms for explaining this segregation still under debate.

Therefore, the present experiment will examine whether negative stimuli (i.e., fearful vs. disgusting) can be differently prioritized in the attentional processing at a neural level occurring “automatically” and whether they could be modulated by HRV influences. Neural responses will be recorded using a high-temporal resolution technique (ERPs) to characterize temporal dynamics of exogenous attention toward emotional distractors during a concurrent but distinct target distractor paradigm. The use of this paradigm where target stimulation (on which voluntary attention should be devoted to accomplish a given task) and task-irrelevant information (emotional distractors) are distinct but appear at the same time has been considered optimal to investigate attentional capture mechanisms [12]. Advanced source estimation tools will also be applied. On these findings we will try to build an explanation of how brain– heart connections may modulate the triggering of exogenous attention to different types of relevant environmental events, specifically when threat-related stimulation appears (fearful vs. disgust).

## Methods

### Subjects

Thirty undergraduate students (14 male) from the Universidad de Jaén, with an age range of 18–26 years old (mean = 21.67, standard deviation = 2.22), voluntarily participated in this experiment. All subjects gave written informed consent and reported normal or corrected-to-normal visual acuity. The Ethics Committee of the Universidad de Jaén approved the study.

### Stimuli and procedure

The experimental paradigm used in this experiment comprised of the presentation of visual stimuli which were composed of two different elements that appeared at the same time on a computer screen: 1) thirty emotional pictures which were used as task-irrelevant stimuli (distractors); and 2) a set of two digits located at the center of each picture being the elements to which participants were asked to voluntarily attend for performing an experimental task (targets). Thus, all participants completed a digit categorization task while they were exposed to three types of distractor pictures: fearful (F), disgusting (D), and emotionally neutral (N). Emotional pictures were selected from two main sources, the International Affective Picture System (IAPS) [35] and Emomadrid [36] on the basis of an independent rating study. Ten pictures conveying fear (IAPS 1525, 1932, 2692, 6190, 6244, 6350, 6510, 9622 and Emomadrid 0578, 0579), disgust (IAPS 9373, 9830 and Emomadrid 0563, 0564, 0569, 0574, 0586 and supplemented with similar pictures found from publicly available sources) and neutral (IAPS 1910, 2512, 2580, 7000 and Emomadrid 0334, 0337, 0338, 0339, 0414, 0584) affective meaning were used. The size for all stimuli was 75.17° (width) and 55.92° (height). Each of these pictures contained two central digits (4.93° height) colored in yellow and outlined in solid black so they were clearly distinguished from the background. As illustrated in Fig. 1, each picture was displayed on the screen for 350 ms and stimulus onset asynchrony was always 3000 ms. The experimental task was exclusively related to the central digits, which could be both even, both odd or one even and the other odd. Participants had to categorize them “as accurately and rapidly as possible”, as congruent or incongruent (in even/odd condition) by pressing different buttons. One key was pressed by the participants if both digits were even or if both were odd (i.e., if they were “congruent”), and a different key if one central digit was even and the other was odd (i.e., if they were “incongruent”). Twenty combinations of digits were composed, half of them being congruent and the other half incongruent. The whole experimental procedure included a total of 120 trials (40 trials for each type of emotional distractor) divided into three blocks (fear, disgust, and neutral). The same combination of digits was repeated across emotional conditions in order to ensure that task difficulty was the same for F, N, and D and were presented in semi-random order into each block of trials in such a way that the congruent or incongruent conditions never appeared more than twice consecutively. The order of the blocks was counterbalanced across subjects. Each picture was presented through a CRT screen four times along the study (twice for both congruent and incongruent trials). Participants were placed in an electrically shielded, sound-attenuated room and they were instructed to continuously look at a fixation mark located in the center of the screen and to blink only after a beep that sounded 1300 ms after each stimulus onset.

**Fig 1.**
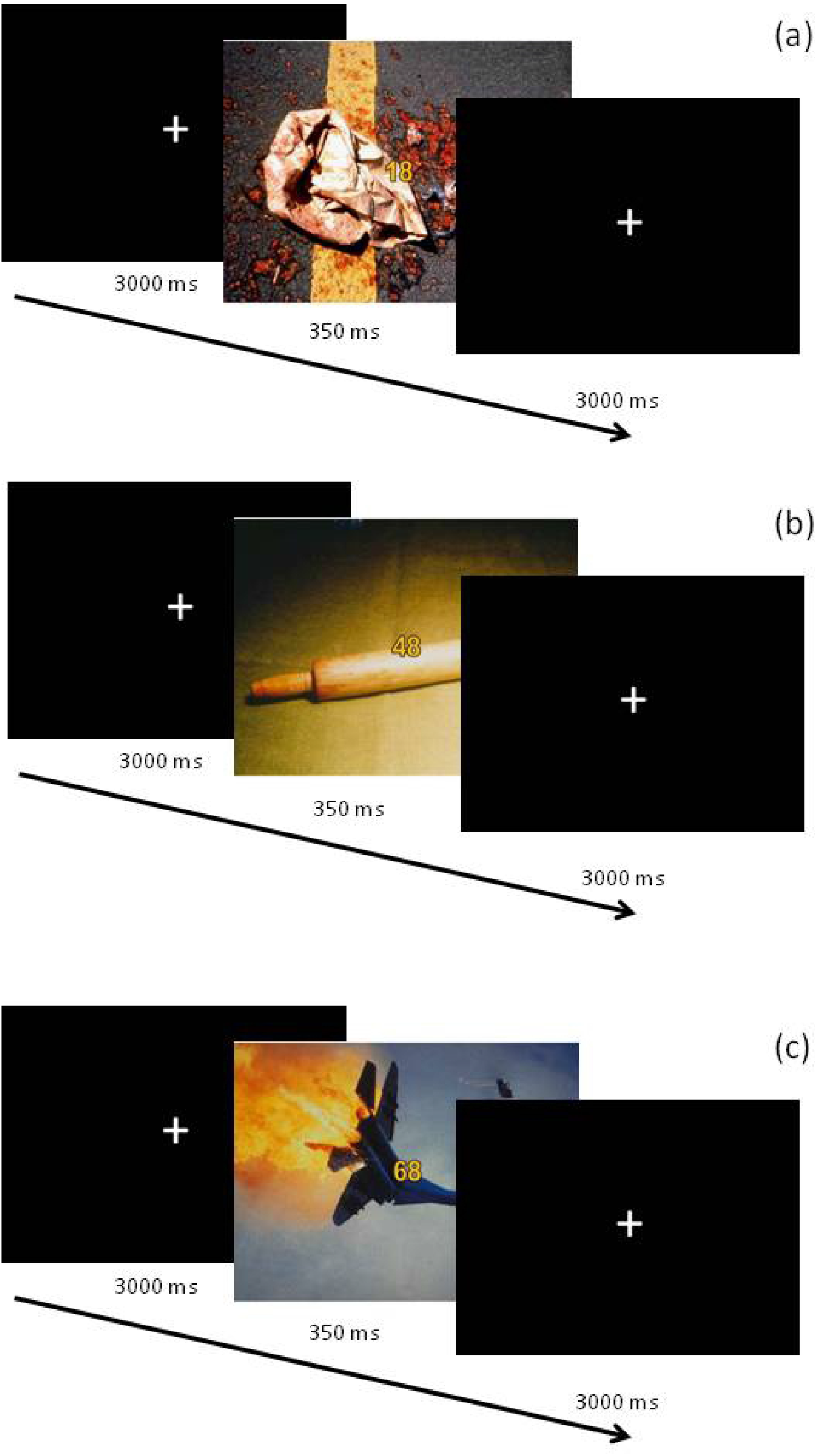
Example of the sequence of events in disgust (a), neutral (b), and fear (c) block of trials.

Images used as distractors were selected from an independent pilot study according to their assessments in valence and arousal (the two main dimensions explaining the principal variance of emotional information), which were similar for fear and disgust categories. Additionally, a subjective criterion was also applied to select pictures representing either fear or disgust images avoiding stimuli combining both emotions.

After the recording session, participants filled out a multidimensional scale for each picture in order to confirm that their assessments on the valence (1, negative to 5, positive), arousal (1, relaxing to 5, arousing), fearfulness (1, minimum to 5, maximum), and disgustingness (1, minimum to 5, maximum) content of the stimulation were those assumed a priori (see Table 1). One-way repeated measured ANOVAs were performed using stimuli (three levels: D, N, and F) as a factor. In order to test the significance of pairwise contrasts, a Bonferroni test was applied. Fearfulness was significantly higher for F than for D stimuli, and disgustingness was significantly higher for D than for F stimuli (p < 0.05 in both cases). Both valence and arousal were higher for F than for D pictures (p < 0.01 in both cases). N stimuli were significantly lower in fearfulness, disgustingness, valence, and arousal than were F and D stimuli (p < 0.01).

**Table 1.** Means and standard deviations (in parenthesis) for the valence, arousal, fearfulness and disgustingness of the emotional distractors (fear, disgust neutral) evaluated by the whole sample of participants. Information about ANOVA p-values for each statistical contrast is included.

|  | Fear | Disgust | Neutral | p-value |
| --- | --- | --- | --- | --- |
| Valence | 2.066 (.501) | 1.761 (.386) | 3.264 (.241) | <.001 |
| Arousal | 4.130 (.378) | 3.466 (.458) | 2.735 (.274) | <.001 |
| Fearfulness | 3.240 (.876) | 1.329 (.335) | 1.040 (.119) | <.001 |
| Disgustingness | 1.316 (.381) | 3.323 (.675) | 1.109 (.146) | <.001 |

### EEG recording and pre-processing

Continuous electroencephalographic (EEG) activity was recorded using an electrode cap (ElectroCap International) with 28 homogeneously distributed scalp electrodes. All electrodes were referenced to the nose tip electrode. Electrooculographic (EOG) data were recorded supra- and infra-orbitally (vertical EOG), as well as from the left versus right orbital rim (horizontal EOG). Electrode impedances were kept below 5 kΩ. An online bandpass filter from 0.3 Hz to 40 KHz was used (3 dB points for −6dB/octave roll-off), and the digitization sampling rate was set to 500 Hz. Off-line pre-processing was performed using Brain Vision Analyzer software (Brain Products). The continuous EEG recording was divided into 900 ms epochs for each trial, beginning 200 ms before stimulus onset. Baseline correction was made using the 200 ms period prior to the onset of the stimulus. Trials in which subjects responded erroneously or did not respond were eliminated. A careful visual inspection was then carried out where epochs with eye movements or blinks were eliminated. This artifact and error rejection led to the average acceptance of 27 disgusting, 27.9 neutral, and 27 fearful distractor trials. The number of trials did not statistically differ among distractor types. The ERP averages were categorized according to each distractor type: fear, disgust and neutral.

### HRV recording and pre-processing

Resting HRV was obtained from the electrocardiogram (ECG) data recorded using three Ag-AgCl electrodes placed following lead II through a Biopac MP100 system in baseline, just before the experimental task and the EEG recording started. The ECG was sampled at 1000 Hz and the beat-to-beat HR data were extracted through Acknowledge 3.9 software. This process yielded a baseline interbeat interval time series of seven minutes duration (longer than the one minute recommended as the minimum necessary according to the Task Force guidelines [37]. Frequency domain analyses were performed on these data using a custom HRV package. Spectral analyses using an autoregressive algorithm following the Task Force guidelines [37] were performed. The frequency domain measure of high frequency (HF: 0.15–0.4) power that has been associated with respiratory-modulated parasympathetic outflow was used to index vagally mediated HRV. Spectral estimates of power (ms^2^) were transformed logarithmically (base 10) to normalize the distribution of scores. Two independent groups of participants were formed on the basis of a median split on their baseline logHF [median = 4.94; low HRV = 3.9 (s.d. = 0.93); high HRV = 5.7 (s.d.= 0.48)] as it has been recommended [38] and extensively used [21, 23, 34, 39].

### Data analysis

#### Behavioral analysis

Mean reaction times (RTs) of correct responses and error rates derived from the task were analyzed. Repeated-measures ANOVAs on each measure were computed using emotional pictures (three levels: F, D, and N) and group of HRV (two levels: high and low) as factors. The Greenhouse–Geisser epsilon correction was applied to adjust the degrees of freedom of the F ratios where necessary, and post hoc comparisons to determine the significance of pairwise contrasts were performed using the Bonferroni procedure (alpha < 0.05). As a measure of effect size, partial η-square (η^2^_p_) is reported for significant effects.

#### ERP analysis

Detection and quantification of ERP components was carried out through covariance-matrix-based temporal principal component analysis (tPCA) explaining most of the brain electrical activity variance. This technique has been strongly recommended for these tasks since its application avoids the subjectivity of selecting “time windows of interest” based on visual inspection for analyses of grand-averaged ERPs. It can lead to several types of misinterpretation, especially when high-density montages are employed (see [40] for a more detailed description of the tPCA procedure and its advantages). The main advantage of tPCA is that it represents each ERP component with its “clean” shape, extracting and quantifying it free of the influences of adjacent or subjacent components [40-41]. Indeed, the waveform recorded at a site on the head over a period of several hundreds of milliseconds represents a complex superposition of different overlapping electrical potentials. Such recordings can stymie visual inspection. In brief, tPCA computes the covariance between all ERP time points, which tends to be high between those time points involved in the same component, and low between those belonging to different components. The solution is therefore a set of independent factors made up of highly covarying time points, which ideally correspond to ERP components. *Temporal factor scores*, the tPCA-derived parameter in which extracted temporal factors may be quantified, is linearly related to amplitude. In the present study, the decision on the number of components to select was based on the scree test [42]. Extracted components were submitted to promax rotation, as recently recommended [43].

As previously mentioned, the analyses required that the ERPs be recorded at 28 globally distributed scalp points. Signal overlapping may also occur at the space domain. At any given time point, several neural processes (and hence, several electrical signals) may concur, and the recording at any scalp location at that moment is the electrical balance of these different neural processes. Thus, ERP components frequently behave differently in some scalp areas to others (e.g., present opposite polarity). While tPCA allows solving temporal overlapping of ERP components, spatial PCA (sPCA) separates ERP components along the space. In this sense, each spatial factor would ideally reflect one of the concurrent neural processes underlying each temporal factor. Therefore, this configuring and quantifying scalp region system is preferable to an a priori subdivision into fixed scalp regions for all components, as sPCA demarcates scalp regions according to the real behavior of each scalp-point recording (basically, each region or spatial factor is formed with scalp points where recordings tend to covary). Consequently, the shape of the sPCA-configured regions is functionally based, and scarcely resembles the shape of the traditional, geometrically configured regions. sPCAs were carried out for each of the temporal factors. This regional grouping was determined through a covariance matrix-based spatial PCA. Moreover, each spatial factor can be quantified through the *spatial factor score*, a single parameter that reflects the amplitude of the whole spatial factor. Additionally, in this case, the decision on the number of factors to extract was based on the scree test. Extracted spatial factors were submitted to promax rotation. This analysis procedure, comprising of both tPCA and sPCA, has been recommended for exploring emotional processing through ERPs [44].

Finally, repeated-measures ANOVAs on temporospatial factor scores were carried out with respect to emotional pictures (three levels: F, D, and N) and HRV group (two levels: high and low). The Greenhouse–Geisser epsilon correction was applied to adjust the degrees of freedom of the F ratios where necessary, and post hoc comparisons to determine the significance of pairwise contrasts were performed using the Bonferroni procedure (alpha < 0.05). Effect sizes were also reported using the partial η-square (η^2^_p_) method.

#### Source estimation analysis

In order to three-dimensionally explore cortical regions that could account for the experimental effects at the scalp level, exact low-resolution brain electro-magnetic tomography (eLORETA) [45-46] was applied to relevant temporal factor scores in accordance with the ANOVA results, as it will be explained later. eLORETA is a 3D, discrete linear solution for the EEG inverse problem [47] which refers to a three-shell spherical model registered to the MNI305 digitized structural human brain atlas template. Solutions are given, therefore, in three coordinates: “x” is the distance in millimeters to the right (+) or left (-) of midline; “y” is the distance anterior (+) or posterior (-) to the anterior commissure; and “z” is the distance above (+) or below (-) a horizontal plane through the anterior and posterior commissures. Although, in general, solutions provided by EEG-based source location algorithms should be interpreted with caution due to their potential error margins, LORETA solutions have shown significant correspondence with those provided by hemodynamic procedures in the same tasks [48-50]. In the present experiment, we tried to minimize this potential error margin using the tPCA-derived factor scores instead of voltages. This strategy has provided more accurate source estimation analyses [2, 51]. In its current version, eLORETA computes the current density at each of 6239 voxels mainly located in the cortical gray matter and the hippocampus of the digitized Montreal Neurological Institute (MNI) standard brain.

Therefore, in order to identify the brain regions underlying exogenous attention processes triggered by emotional stimulation, three-dimensional current density estimates for relevant temporal factor scores that were sensitive to experimental manipulations according to ERP results were computed for each participant and each experimental condition. Subsequently, the voxel-based whole-brain eLORETA-images (6239 voxels) were compared among three experimental conditions (F, D, and N: emotional distractors). To that aim, eLORETA built-in voxelwise randomization tests (5000 permutations) based on a statistical non-parametric mapping methodology were used. As explained by Nichols and Holmes [52], the non-parametric methodology inherently avoids multiple comparison-derived problems and does not require any assumption of normality. Voxels that showed significant differences between conditions (log-F-ratio statistic, two-tailed corrected p < 0.05) were located in anatomical regions and Brodmann areas (BAs). In order to explore exogenous attentional processing related to the three types of emotional distractors (three levels: F, D, and N) and the two groups of HRV (two levels: high and low), current densities of different regions of interest (ROIs; radius = 5 mm) were submitted to ANOVAs.

## Results

### Behavioral results

Average values for RTs and error rates for each emotional distractor are shown in Table 2. Two separate repeated-measures ANOVAs were conducted on RTs and error rates including emotional distractor (three levels: F, D, and N) and HRV group (two levels: high and low) as factors. Although both RTs and error rates for D stimuli were slowest and highest, as it can be observed in Fig. 2, a significant main effect of emotional distractor was only found for error rates [F(1.91,53.53) = 3.297, p = .047, η^2^ = 0.105] but not for RTs [F(1.82,50.85) = 3.075, p = .06]. Disgusting distractors provoked higher error rates than neutral distractors (p < 0.05). There was no significant main effect for HRV group (RTs [F(1,28) = 2.413, p = .132]; error rates [F(1,28) = .093, p = .763]). Finally, the interaction between emotional distractor and HRV was significant for RTs [F(1.82,50.85) = 5.419, p = .009, η^2^ = .162] but not for error rates [F(1.91,53.53) = .336, p = .706]. Posterior analysis conducted on this interaction revealed slower RTs to D distractors compared to N (p = .01) and F stimuli (p = .04) in the high HRV group. Additionally, low HRV group showed faster RTs to F distractors compared to D (p = .048) and N (p = .01).

**Table 2.**
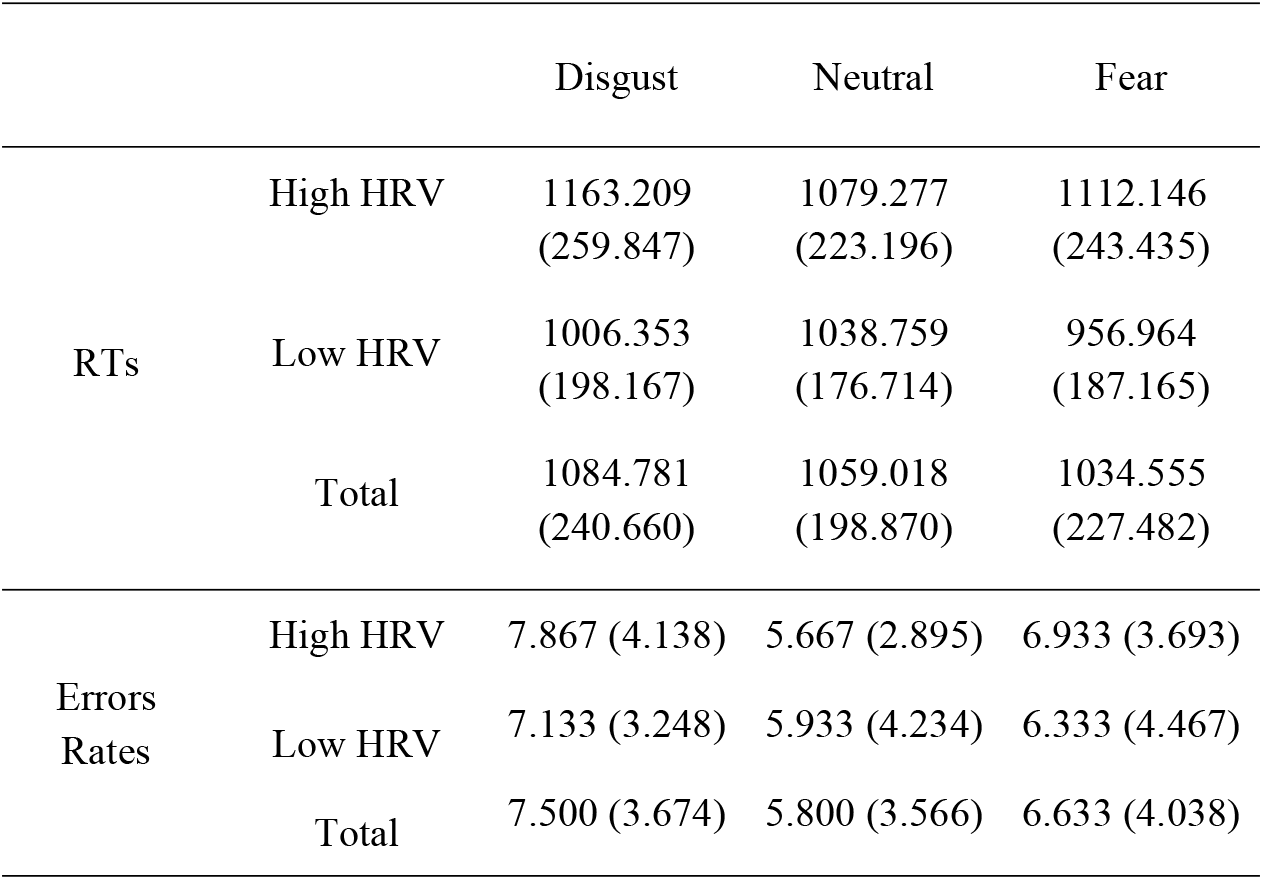
Means and standard deviations (in parenthesis) of RTs and errors rates to disgust, neutral and fearful distractors.

**Fig 2.**
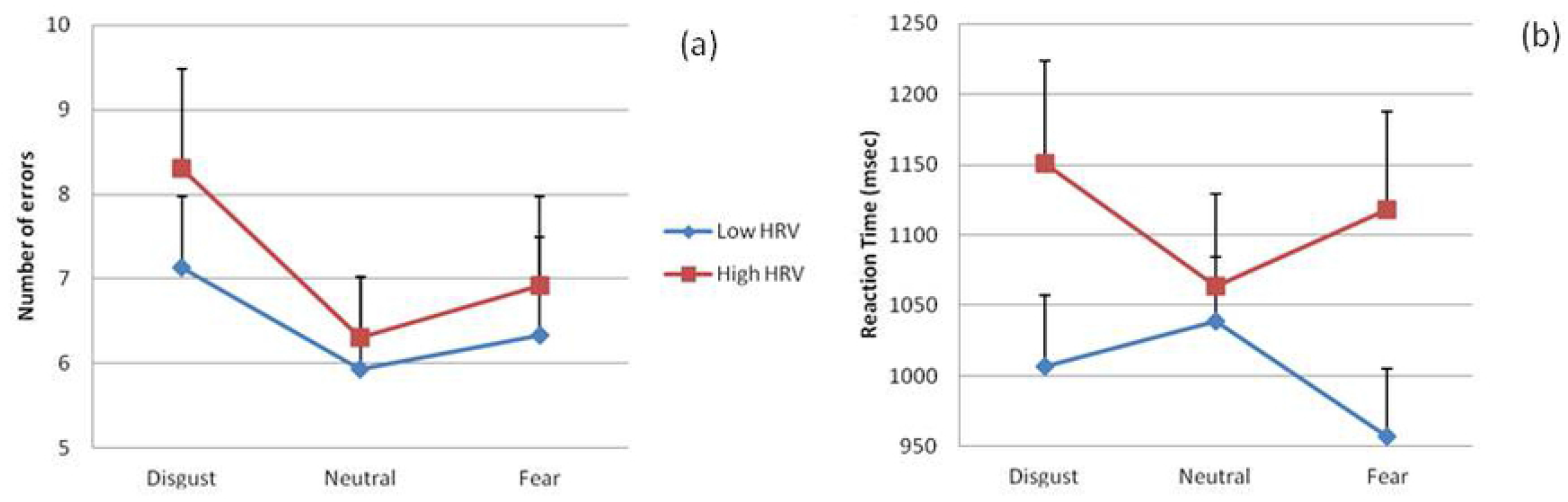
Number of errors (a) and reaction times (b) related to the different emotional distractors in each HRV group. Error bars = standard errors.

### ERP results

Grand averages related to exogenous attentional brain responses for the three emotional distractors, once the baseline value (pre-stimulus recording) has been subtracted from each ERP, are displayed in Fig. 3. This figure mainly shows ERP activity (P1, P2, and N2) where the most relevant experimental effects related to attentional capture processes are clearly appreciable. These effects will be subsequently detailed.

**Fig. 3.**
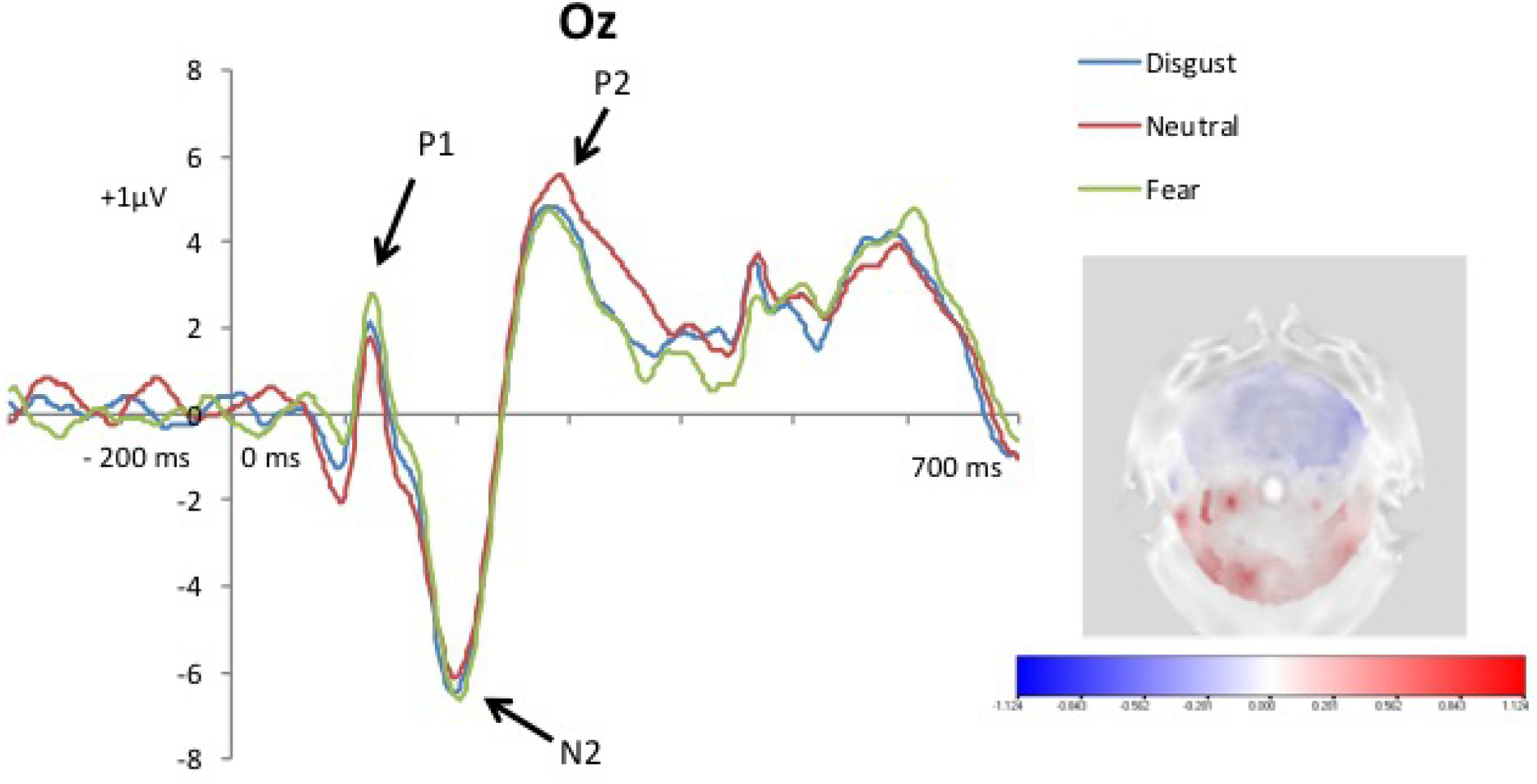
Grand averages related to exogenous attentional ERP responses (P1, P2 and N2) for the three emotional distractors.

As a consequence of the tPCA application, eight temporal factors (TFs) were extracted from the ERPs (see Fig. 4). Extracted factors explained 83.79% of the total variance. According to the factor peak latency and topography distribution, TF8 and TF6 (peaking at 138 and 228 ms, respectively) were identified at fronto-central and centro-parietal sites of the scalp and associated with ERP components signaled in the grand averages as P1 and P2 (see Fig. 3). Furthermore, TF7 (peaking at 176 ms), maximal at frontal scalp sites, was related to N2 (Figs 3 and 4).

**Fig. 4.**
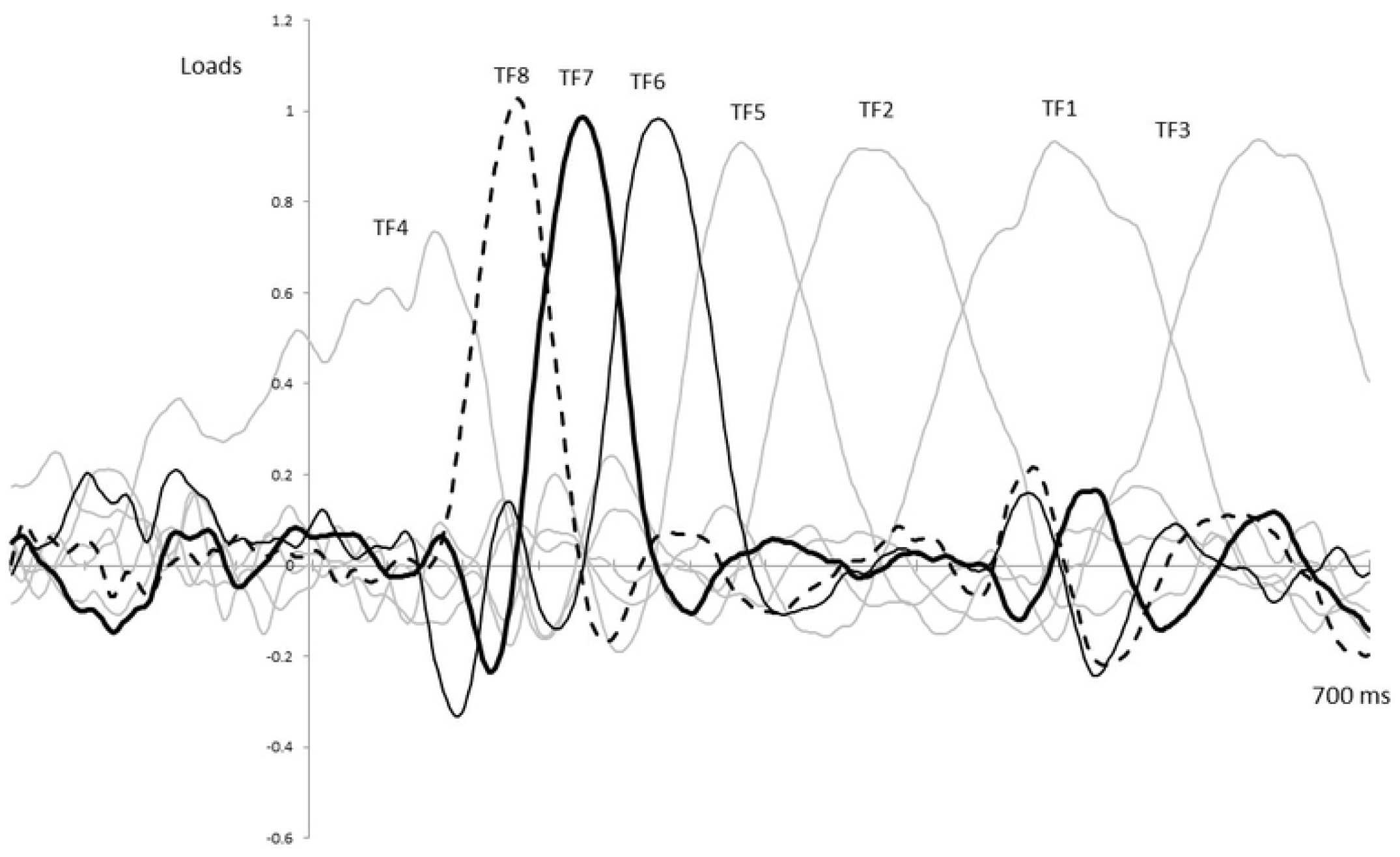
Temporal factors extracted through the application of the tPCA.

Subsequently, the sPCA extracted three spatial factors for each temporal factor, except for TF5 and TF6, where only two spatial factors were extracted. Therefore, the temporospatial PCA yielded a total of 22 factor combinations. Repeated-measures ANOVAs on each temporospatial factor were carried out for exploring exogenous attention (i.e., P1, P2, N2 at different scalp areas) triggered by emotional distractors (three levels: F, D, and N) with respect to the HRV group (two levels: high and low).

Table 3 also provides the statistical details of these analyses including main effects of emotional distractor and the interaction effects between emotional distractor by HRV group. As can be appreciated in this table, a clear effect related to the emotional content conveyed by distractor pictures was revealed for both P1 and P2 components. Specifically, posterior scalp regions of P1 (corresponding to SF1 and SF2) showed a significant main effect of emotional distractor [SF1: F(1.97,55.26) = 5.729, p = .006; η^2^ = 0.170; SF2: F(1.95,54.65) = 3.429, p = .041; η^2^ = 0.109], with higher amplitudes for trials including fearful distractors than neutral ones (p < .05). The main effect of emotional distractor also reached significant results for P2 [SF2: F(1.99,55.696) = 3.710, p = .031; η^2^ = 0.117], showing a complementary pattern in which brain amplitudes at frontal regions were higher for disgusting compared to neutral distractors (p = .050). However, no interaction effects involving HRV were found for these ERP components.

**Table 3.**
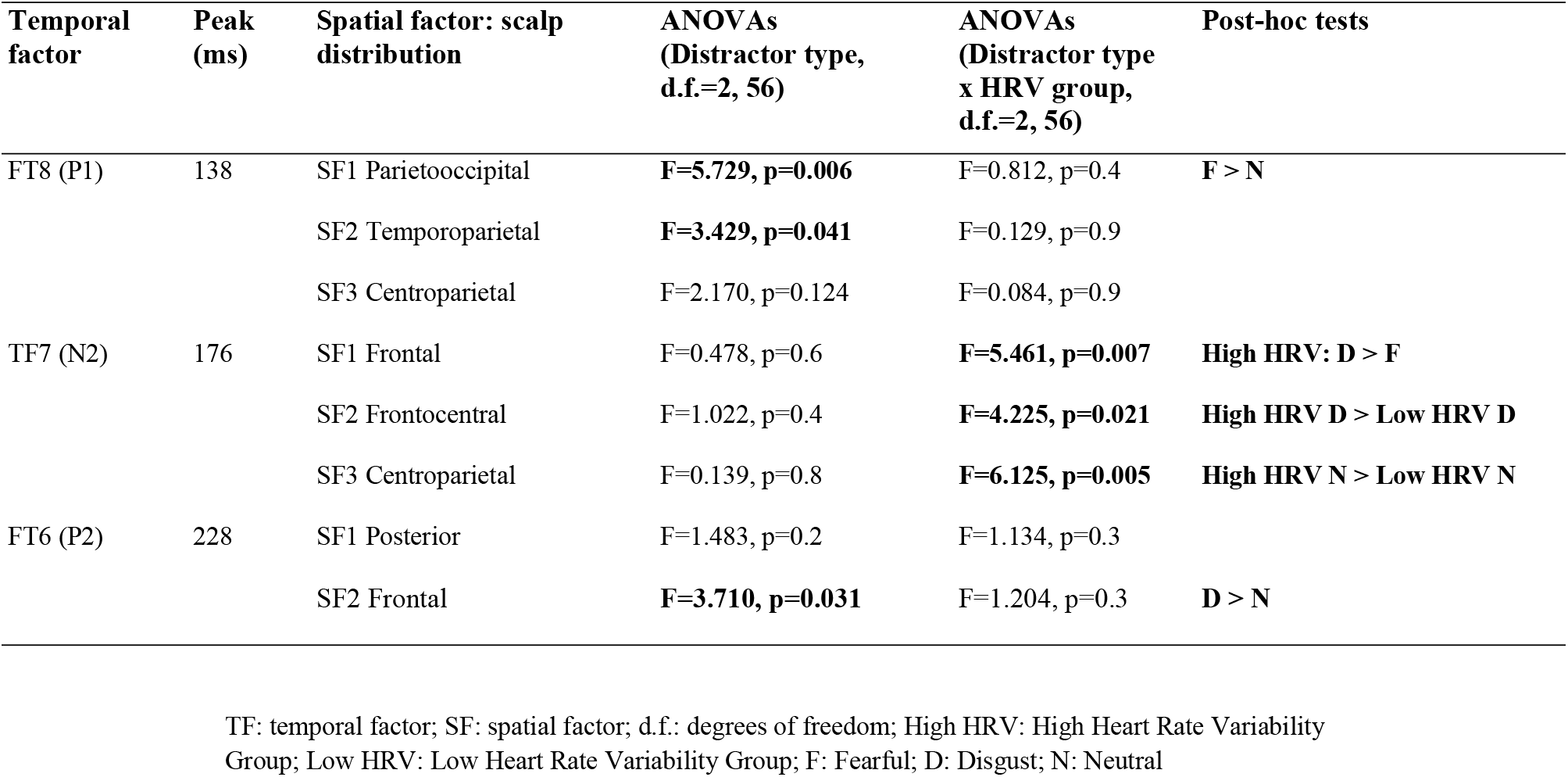
Description and statistical results for the temporal factors sensitive to experimental manipulations extracted by temporospatial Principal Component Analysis. Post-hoc results are shared by all spatial factor belonging to each temporal factor.

Finally, with respect to N2, different scalp regions reached statistical significance for the interaction between emotional distractor by HRV group [SF1: F(1.97,55.18) = 5.462, p = .007; η^2^ = 0.163; SF2: F(1.9,53.27) = 4.225, p = .021; η^2^ = 0.131; SF3: F(1.89,52.90) = 6.125, p = .005; η^2^ = 0.179]. Post hoc tests showed enhanced N2 amplitudes for disgusting compared to fearful emotional distractors. This pattern was only exhibited for the high HRV group (p < 0.01; see Figs. 5a and 5b). Although the N2 amplitude was different between fearful compared to neutral distractors, it did not reveal significant differences for any scalp regions (p = 1). Furthermore, high HRV subjects exhibited frontal N2 augmented amplitudes for both disgust and neutral distractors compared to low HRV group (p < 0.05). No other significant data were found on the rest of the ERP components with respect to this interaction effect. Finally, a main effect of HRV group (F(1,28) = 4.294, p = .048, eta = .133) was found for N2, which presented a higher amplitude in high HRV than low HRV individuals. Mainly both frontal and central regions were sensitive to this comparison.

**Fig. 5.**
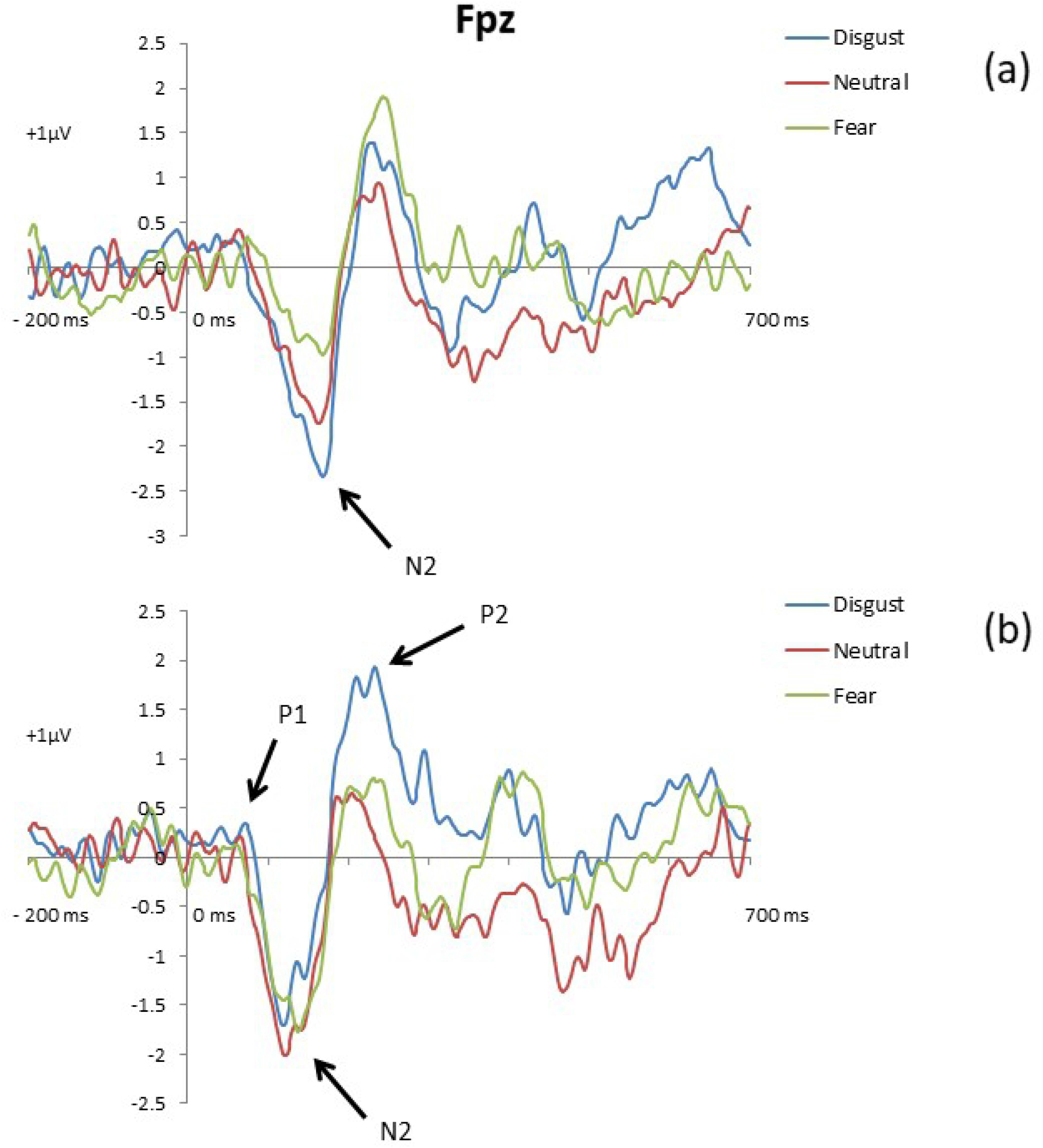
ERP amplitudes to the three emotional distractors in high (a) and low (b) HRV groups at FPz.

### Source estimation results on N2

In order to three-dimensionally estimate cortical regions that could account for the experimental effects observed in N2 regarding the interaction between emotional distractor by HRV group, a final analytical step was computed. To achieve this, N2 TF scores of each subject, electrode, and condition were submitted to eLORETA. Then, the voxel-based whole-brain eLORETA-images (6329 voxels) were compared among the three emotional distractor conditions for the whole sample of participants. To this aim, statistical non-parametric randomization tests were performed. According to the computed comparisons related to N2 activation, disgusting distractors were associated with enhanced activity in several voxels within BA13 (insula; MNI coordinates: x = 35, y = 5, z = 15) for high HRV compared to low HRV subjects in response to disgusting distractors (see Fig. 6). Activation differences among the remainder of the distractor conditions did not reach significance. No other ROIs were sensitive regarding this interaction between emotional distractor by HRV group.

**Fig. 6.**
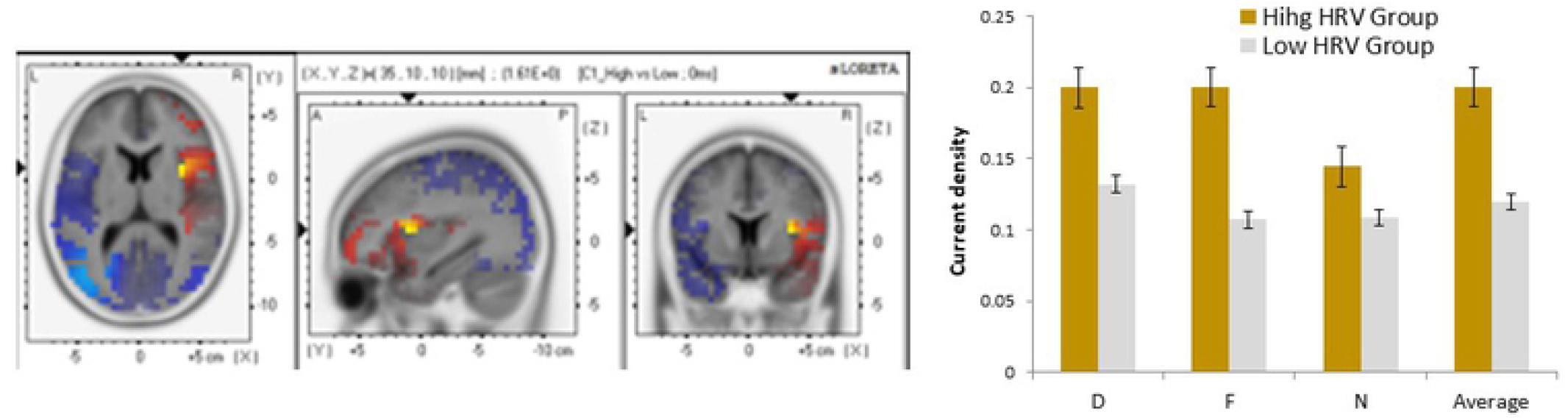
Insula activation to disgusting distractors (low vs. high HRV). Average current densities for each condition are also shown. (D, disgusting distractors; F, fear-evoking distractors; N, neutral distractors). Error bars represent standard errors.

## Discussion

The current study points out that to understand the differential effect of fearful and disgusting stimulation on exogenous attention it is important to consider the time course of its different subprocesses. Fear seems to be processed at a very early stage such as 100 ms after stimulus onset, producing a rapid response in the whole sample of participants. Disgust would also capture exogenous attention at a very early phase, although later than fear. In fact, disgusting distractors attracted exogenous attention resources more efficiently than fear at 200 ms after stimulus onset, as reflected by its effects on the N2 and P2 components of the ERPs. Interestingly, the augmented N2 response to disgust distractors was only present in those participants having a high HRV level.

These apparently puzzling results can well be understood from an evolutionary perspective, which is in agreement with previous proposals describing that exogenous attentional involves different functional phases or processes (see a review in [12]). Prior studies have distinguished different ERP components (peak latency from 100 to 250 ms) in response to visual stimulation reflecting the temporal dynamics of exogenous attention towards emotional events (e.g., [2, 5]). Early functional phases or processes of exogenous attention would be related to a greater mobilization of processing resources toward negatively valenced stimuli indicating probably a negativity bias, which would have adaptive and evolutionary advantages for survival (see reviews, [12, 53]). This effect would confirm the biological importance of emotional stimuli compared to neutral [4, 6, 54-55]. On the contrary, during later phases, attentional resources would be directed preferentially to positive and neutral events that do not require such an urgent response as negative stimuli do [55]. However, a direct comparison between present data and this previous evidence cannot be made for different reasons: 1) experimental paradigms were different; 2) we did not include positive stimuli; 3) in previous investigations negative pictures used depicted only fear-related but not disgust-related content. Even though, our results are not exclusive with those reported by other studies, but rather are complementary.

According to the statistical results, fearful distractors elicited the largest posterior P1 amplitudes. It has been indicated that fear eliciting stimulation signals an imminent threat that calls for an immediate and fast response while disgust stimuli do not. In support of this idea, previous studies on exogenous attention have reported enhanced parieto-occipital P1 amplitudes to fear evoking stimuli used as distractors [3, 56-57]. In the same line, attentional effects to disgust stimuli have not been reflected in P1 amplitude, which has been interpreted as its effects occur later [58]. Different findings converge to propose that higher P1 amplitudes would reflect amplification of sensory processing (as a subprocess belonging to exogenous attention) to fear stimulation gaining access for awareness and processing it to a deeper extent [5]. This enhancement of attentional resources to fearful events would involve a clear adaptative advantage in biological terms, playing a crucial role for survival [1]. In this sense, sensory visual cortices would increase their neural activity mediated by top–down attention mechanisms where both the amygdala and ventral prefrontal regions would generate rapid saliency signals for fear stimulation [59-61].

On the other hand, ERP components peaking around 200 ms showed an increase of attention resources for processing disgusting distractors. Specifically, P2 component was characterized by two spatial components (frontal and posterior) through spatial principal component analyses. Although the posterior P2 wave showed no modulations regarding exogenous attention, disgust distractors elicited larger anterior P2 amplitudes than neutral distractors. Furthermore, disgust distractors produced larger N2 amplitudes than fearful stimuli, but this modulation was only true for the high HRV group. Several investigations have found similar modulations in P2 showing greater sensitivity toward negative events, considering this component as a neural index of exogenous attention [55, 62-63]. Prior studies have highlighted that disgust is a kind of event capable of attracting automatic attention resources even more efficiently than fear [14]. Although this effect was also found around 200 ms after stimulus onset, N2 modulations was not previously reported and possible influences of HRV on neural responses linked to exogenous attention has not been described yet.

The selective enhancement of exogenous attention to negatively valenced distractors (i.e, fearful and disgusting) deserves further consideration. When the organism is involved in the performance of a given task and a potentially threatening distractor appears, the most adaptive response is to do a quick evaluation to explore whether such new stimulus seems to be dangerous in order to initiate a fight or flight response [1]. However, in the case that the outcome of such initial evaluation does not indicate an imminent danger, but the stimulus includes ambiguous information, as happens with disgusting stimuli that can bring death (e.g., illness) or life (e.g., edible food), the most adaptive response would be mobilize attentional resources to analyze it more deeply. Our findings suggest that this attentional adaptive response is evident for high HRV participants as shown by both enhanced N2 amplitudes and faster RTs to disgusting distractors compared to fearful ones. Also disgusting provoked higher error rates than neutral distractors. N2 family deflections have been described as emotion-sensitive ERP waves showing enhanced allocation of exogenous attention resources toward negative stimulation compared to neutral ones [62, 64]. In a previous study, we also found that disgusting distractors capture attention more efficiently than fearful distractors [14] as reflected by neural responses peaking around 200 ms after stimulus onset. Other studies have offered evidence indirectly supporting the suggestion that disgusting distractors capture exogenous attention with latency responses around 200 ms. Van Hooff et al [11] manipulated the temporal interval between cue onset (disgust, fear, or neutral picture) and target showing that targets presented 200 ms after disgust cue onset were identified less accurately and more slowly than targets presented after fear or neutral cues, indicating a more efficient capture of attention by disgust than by fear cues. Similarly, other studies focused on the study on automatic attention to negative stimulation (disgust and fear), have described longer RTs to targets that appeared 200 ms after the disgusting pictures [65]. Although ERPs were not recorded and the methodology used in their study does not allow exploration of the time course of the differential responses to fear and disgust, it has been suggested that attentional response for fearful pictures seems to occur earlier and more automatically than for disgusting events, since just the quick registration of their rough content is enough to trigger the appropriate fight or flight reaction [65].

The influence of HRV on the very early phases of the exogenous attentional processing (i.e., P1 and P2 ERP deflections) was not observed. However, an interesting effect was evident in a later phase, as reflected by the augmentation of N2 amplitudes to disgust distractors, only for the high HRV group. Our results also indicated that this enhancement of N2 seem to be related to neural activity within the insula. These effect involving participants characterized by a high level of HRV was also prominent at the behavioral level. RTs for trials including disgust distractors were longer compared to neutral and fear ones. No effect of HRV was detected in the error rates.

This pattern of results regarding the role of HRV can also be interpreted from an evolutionary perspective. Following present results, the HRV may play an important role in a late stage of attentional processing, where individuals (i.e., high HRV group) become involved into a more detailed exploration of disgusting stimuli, as reflected by RTs and N2. However, low HRV participants did not show the same attentional pattern. They seem not to be able to disengage their initial attentional response to fear distractors as manifested by their short RT responses, which is typical of hypervigilance responses [34]. Similar results have previously been reported by our group. A deeper cardiac deceleration was found to disgusting distractors compared to fear and neutral ones in an exogenous attention task only in high HRV participants. The low HRV group showed no differences in cardiac deceleration to the three types of emotional distractors [23]. However, early attentional responses to threat are required to face with this kind of stimulation for survival. In this case, higher neural responses were detected in P1when fearful distractors appeared across all participants, regardless of their HRV level.

As it mentioned, neural activity within the insula was related to the enhanced N2 during disgust distractor trials for high HRV individuals. The insula has been associated with the processing of negative emotional stimuli [12, 66-67], especially with the emotion of disgust [13] as well as with interoception and bodily signals’ processing [68-69]. Importantly, recently the insula has also been suggested as a key structure in two relevant models for the interpretation of our findings. On one side, Carretié [12] included the insula as one of the three main brain areas (together with the amygdala and ventral prefrontal cortex) responsible for the first phase of exogenous attention, called “preattention”. During this phase, a fast evaluation of the environment based on low-level stimulus features is conducted in order to detect relevant stimulation and to trigger reorienting processes. Also, Shafer et al [70] included the insula between those brain areas that reflect automaticity in the emotional processing. On the other side, the relationship between HRV and cerebral blood flow responses elicited by different kind of emotional states such as happiness, sadness, disgust neutral has been previously explored [24]. Their results showed an association between HRV and activity in the insula, but the specific role of each specific emotion on this heart and brain link was not explored. Smith et al [71] presented a version of the Neurovisceral Integration Model in which the nervous system structures that contribute to vagal control are organized in an eight-level hierarchy, increasing the range and complexity of function at each level. The insula plays a primary role at level six, where the structures implicated in the vagal regulation based on perceptual representation of one’s current somatic/visceral state are included. Taking into consideration the role of the insula indicated by these authors from different perspectives, it could be suggested that this structure may be contributing to the preattention phase by offering important information about the current bodily state that may help to regulate vagal control as well as the attentional responses.

Vagally mediated HRV has been structurally and functionally linked to emotional regulation. It has been argued that prefrontal, anterior cingulated, and insula cortices constitutes a neural network with bidirectional communication with the amygdala and other structures implicated in emotional regulation as well as autonomic regulation of the heart [25]. In this way, HRV becomes an index of emotion, cognition, and health related physiological processes. Regarding emotional regulation, it has been found that high HRV is associated with context appropriate emotional responses, for example, under a startle emotional modulation paradigm [22] or to phasic HR responses in addition to behavioral and self-reported emotional responses [23, 39, 72-73]. Low HRV, however, has previously been related to poor emotional regulation, hypervigilance, and exaggerated responses to innocuous stimuli [21-22, 31], This maladaptive pattern of responses is consistent with faster RTs shown to trials containing fearful distractors and the absence of a P2 response related to disgusting stimuli in the present study in the low HRV participants. Although resting HRV can be understood as a trait like measure, it is not included in the characteristics that have been repeatedly associated with negativity bias to negative stimuli. However, the findings of the present study suggest that HRV may be a more informative index of underlying processes of exogenous attention to emotional stimuli than trait anxiety or disgust sensitivity (see S1 Supporting Information), which posits as a potential tool that may help to understand and disentangle inconsistencies in the previous literature.

Although exposed findings offer relevant evidence on the brain dynamics related to attentional capture by discrete emotions, some potential limitations need to be addressed. In this sense, valence and arousal scores were significantly more negative and arousing for fearful than disgusting distractors. Such differences in these dimensional affective characteristics between emotional stimuli (distinct from their fearfulness and disgustingness, respectively) might have some unexpected influence on attentional capture processes leading to misunderstandings. Nevertheless, although discrete emotions (as a part of affective experience) could be represented in a dimensional space [74], they have their own and intrinsic emotional meaning that makes them unique, apart from the other features. In the current context, future studies combining both discrete and dimensional approaches for representing emotional states are needed to extend knowledge on the capability of emotional stimuli for attracting automatic attention resources. On the other hand, the relatively small sample size used in the present experiment could affect statistical power. To check if our results had the necessary and significant statistical power, we carried out a series of post-hoc power analysis (see S2 Supporting Information). These analyses yielded a high statistical power for the interaction effect between the type of Distractor x HRV group for each of ERP component sensitive to the experimental manipulations related to attentional capture.

In sum, findings derived from the present investigation provide new data about how exogenous attention is captured by different types of negatively valance stimuli. A different time course for fear and disgust is shown by electrophysiological indices. Fear distractors capture automatic attention at very early processing phases (i.e., greatest P1 amplitudes) followed by an augmented modulation in the P2 component in response to disgust events. Data suggest that HRV might contribute to modulate the allocation of exogenous attentional resources for disgusting stimulation as reflected in the N2 amplitudes. Enhanced activation within the insula lead to think that this region may have a relevant role in the early stages of exogenous attention. Future research should be done to better characterize the temporal course of exogenous attention to emotional stimuli, taking into account measures of HRV as a variable that might be important when attention is automatically triggered by potential threats.

## Acknowledgements

The authors would like to thank all participants for taking part in the experiment.

## Author Contributions

Conceptualization: Elisabeth Ruiz-Padial.

Data Curation: Elisabeth Ruiz-Padial.

Formal Analysis: Francisco Mercado

Funding Acquisition: Elisabeth Ruiz-Padial, Francisco Mercado Investigation: Elisabeth Ruiz-Padial.

Methodology: Elisabeth Ruiz-Padial, Francisco Mercado

Project Administration: Elisabeth Ruiz-Padial

Resources: Elisabeth Ruiz-Padial

Software: Elisabeth Ruiz-Padial, Francisco Mercado

Supervision: Elisabeth Ruiz-Padial

Validation: Elisabeth Ruiz-Padial, Francisco Mercado

Writing-Original Draft Preparation: Elisabeth Ruiz-Padial, Francisco Mercado Writing-Review & Editing: Elisabeth Ruiz-Padial, Francisco Mercado.

